# Molecular origin of somatostatin-positive neuron vulnerability

**DOI:** 10.1101/2021.02.16.431515

**Authors:** Toshifumi Tomoda, Akiko Sumitomo, Dwight Newton, Etienne Sibille

**Author notes:** Correspondence to: T. Tomoda; E. Sibille), Etienne Sibille, Ph.D., Centre for Addiction and Mental Health, 250 College Street, Room 134, Toronto, Ontario M5T 1R8, Canada. x. 36582.

## Abstract

Reduced somatostatin (SST) and SST-positive (SST^+^) neurons are hallmarks of neurological disorders and associated with mood disturbances, but their origin are unknown. Chronic psychosocial stress induces behavioral emotionality deficits and deregulates unfolded protein response (UPR) of the endoplasmic reticulum (ER) preferentially in SST^+^ neurons. Here we confirm that chronic stress increases ER stress levels in SST^+^ neurons of mouse prefrontal cortex, and show that genetically suppressing ER stress in SST^+^ neurons, but not in pyramidal neurons, normalized psychosocial stress-induced behavioral emotionality. Forced expression of SST precursor protein (preproSST), mimicking psychosocial stress-induced early proteomic changes, induces ER stress, whereas mature SST or processing-incompetent preproSST does not. Biochemical analyses further show that psychosocial stress induces SST protein aggregation under elevated ER stress conditions. These results demonstrate that SST processing is a SST^+^ neuron-intrinsic vulnerability factor under conditions of sustained or over-activated UPR in the ER, hence negatively impacting SST^+^ neuron functions.

## Introduction

Somatostatin (SST) is a multi-functional neuropeptide expressed in a major subtype of γ-aminobutyric acid (GABA) neurons (Epelbaum et al., 2009). SST-positive (SST^+^) neurons mediate inhibitory neurotransmission mainly in dendrites of excitatory pyramidal neurons (Hendry et al., 1984; Melchitzky and Lewis, 2008; Xu et al., 2010). Preclinical studies demonstrate that SST^+^ neurons regulate information processing at the cortical microcircuit level, and that low SST or inhibiting SST^+^ neuronal functions in corticolimbic brain areas is sufficient to cause excitation-inhibition (E/I) imbalance contributing to mood disturbances (Soumier and Sibille, 2014; Lin and Sibille, 2015; Fee et al., 2021). Low SST levels are reported in a diverse array of brain disorders, including Alzheimer’s disease (AD), Parkinson’s disease (PD), schizophrenia, bipolar disorder, major depressive disorder (MDD) and during normal aging (Martel et al., 2012; Fee et al., 2017), suggesting a common pathophysiological mechanism across deregulated brain conditions (Prévôt and Sibille, 2021).

Chronic psychosocial stress causes alteration of neurocircuitries responsible for controlling behavioral emotionality, in particular those involving hypothalamus and corticolimbic areas of the brain (Willner, 2016). An array of stress paradigms leads to increased corticosterone levels, which trigger rapid SST release from SST^+^ neurons (Epelbaum et al., 2009). SST^+^ neurons are engaged in stress response and particularly vulnerable under chronic stress conditions, including brain conditions such as MDD (Tripp et al., 2011; Guilloux et al., 2012). Using mice exposed to unpredictable chronic mild stress (UCMS), a paradigm that mimics chronic psychosocial stress and recapitulates various neurobiological and behavioral processes implicated in MDD (Willner, 2016), we and others confirmed reduced SST in corticolimbic areas and reported additional genes selectively altered in SST^+^ neurons, notably the eukaryotic translation initiation factor (EIF2A) signaling pathway genes responsible for protein synthesis (Lin and Sibille, 2015, Girgenti et al., 2019). This suggests a SST^+^ neuron-selective deficit of proteomic regulation or proteostasis associated with chronic stress.

EIF2A signaling is a ubiquitous housekeeping mechanism that regulates protein translation and subsequent processing through the endoplasmic reticulum (ER), a major site of posttranslational modification in all cell types (Ron and Harding, 2012). Upon acute cellular stress, increasing levels of misfolded/damaged proteins in the ER can cause ER stress and initiate the unfolded protein response (UPR). This is in part achieved by suppressing EIF2A signaling via phosphorylation of EIF2α, which blocks protein translation as a protective cellular mechanism (Metcalf et al., 2020). However, sustained or overactivated ER stress can induce apoptosis signaling, as reported for AD or other neurodegenerative disorders associated with deficient or misfolded proteins accumulation. For example, pharmacologically suppressing ER stress by inhibiting the protein kinase RNA-like ER kinase (PERK), one arm of the UPR pathway responsible for EIF2α phosphorylation, could paradoxically mitigate neuronal cell death in an AD mouse model (Rozpedek et al., 2015; Gerakis and Hetz, 2018). Given the chronic nature of MDD pathobiology and by analogy to sustained ER stress in AD, we previously tested the efficacy of a PERK inhibiting compound in UCMS mice and showed that it could mitigate chronic stress-induced behavioral emotionality (Lin and Sibille, 2015), suggesting that overactivated ER stress may serve as a pathophysiological mechanism underlying MDD.

While accumulating evidence highlights the role of chronic psychosocial stress in low SST and selective SST^+^ neuron deficits (Fee et al., 2017), underlying mechanisms are missing. For instance, direct evidence linking a putative causal role of ER stress to the SST-related pathology is lacking and the molecular mechanisms linking overactivated ER stress/UPR and SST^+^ neuron vulnerability are unknown. Whereas deficient or misfolded proteins accumulation induce ER stress in neurodegenerative disorders (Hetz and Saxena, 2017), evidence also suggest that excessive processing of normal endogenous proteins can induce ER stress. For instance, in diabetes, chronic elevated demand for insulin production and secretion through the ER and trans-Golgi network causes exacerbated ER stress in pancreatic β cells (Arunagiri et al., 2018). Given that SST is a stress-inducible neuropeptide (Negro-Vilar and Saavedra, 1980; Arancibia et al., 1984, 2000; Chen and Du, 2002; Priego et al., 2005; Polkowska and Wankowska, 2010; Prévôt et al., 2017) that is produced as a precursor form (preproSST) and processed through the ER to become a mature bioactive peptide in SST^+^ neurons (Goodman et al., 1983), we tested the hypothesis that (i) chronic stress may cause elevated ER stress in SST^+^ neurons through increased need for production and processing of preproSST, and that (ii) as a result of overactivated ER stress/UPR, SST^+^ neuron-intrinsic factors may undergo compromised proteostasis and SST aggregation, together contributing to low free SST and SST^+^ neuron dysfunctions.

## Results

### Transcriptomic evidence for SST^+^ neuron vulnerability following unpredictable chronic mild stress (UCMS) in mice

Our previous transcriptome analysis in corticolimbic areas of UCMS mice identified selective downregulation of EIF2A pathway genes in SST^+^ neurons, and suggested chronic stress-induced SST^+^ neuron deficits through elevated ER stress (Lin and Sibille, 2015). To further investigate the degree of cell type-selective vulnerability and potential underlying mechanisms, we re-analyzed data from a single cell-type transcriptome study in mice exposed to UCMS, in which 4 major neuronal subtypes (i.e., pyramidal, SST^+^, parvalbumin (PV)^+^, or vasoactive intestinal peptide (VIP)^+^ neurons) in medial prefrontal cortex (PFC) were sampled via laser capture microdissection and subjected to RNAseq (Newton et al., 2020). Using gene set enrichment analysis (GSEA) of all gene sets related to the ER stress/UPR machinery (i.e., gene ontology (GO) terms containing ER stress, unfolded/misfolded proteins, molecular chaperone, heat shock proteins/response to heat), 2 and 8 GO terms were significantly enriched in pyramidal and SST^+^ neurons respectively, whereas none was enriched in PV^+^ or VIP^+^ neurons (**Fig.S1**). Among the three known arms of ER stress/UPR pathway regulators (i.e. PERK, IRE1, ATF6) (Metcalf et al., 2020), PERK pathway showed the highest enrichment score in SST^+^ neurons, whereas IRE1 and ATF6 showed negative or no enrichment (**Fig.S2A,B**). Analysis of the effects of UCMS on representative genes in each pathway further showed a biased elevation of PERK pathway gene expression, over IRE1 and ATF6 pathways, in SST^+^ neurons (**Fig.S2C**).

This data provides evidence that SST^+^ neurons are enriched in PERK pathway gene expression over other ER stress/UPR regulators and that chronic stress preferentially affects the PERK pathway in SST^+^ cells. This is consistent with the previous findings that SST^+^ cells are vulnerable to chronic psychosocial stress and that a PERK inhibiting compound suppressed the increase in behavioral emotionality of mice exposed to chronic stress (Lin and Sibille, 2015), together supporting the PERK pathway as the best candidate for further study of the selective vulnerability of SST^+^ cells.

### Elevated ER stress in SST^+^ neurons is sufficient to cause elevated behavioral emotionality

We next validated the elevated ER stress levels via immunofluorescence analysis in UCMS-exposed mice. During 1, 3 and 5 weeks of UCMS, a time-dependent progressive increase in phospho-eIF2α (Ser-51) levels was observed in SST^+^ neurons (labeled green) of PFC equally in male and female mice (*Perk*^+/+^;*Ai6*^lsl-ZsGreen/+^;*Sst*^IRES-Cre/+^; **Fig.1A**; **1B**, black dots). Among 4 known integrated stress response kinases responsible for eIF2α phosphorylation, PERK is the only ER resident kinase. To examine the extent of eIF2α phosphorylation elicited by PERK in UCMS mice, we generated mice with reduced *Perk* gene levels in SST^+^ neurons (*Perk*^flox/+^;*Ai6*^lsl-ZsGreen/+^;*Sst*^IRES-Cre/+^ and *Perk*^flox/flox^;*Ai6*^|s|-ZsGreen/+^;*Sst*^IRES-Cre/+^). The UCMS-induced increase in phospho-eIF2α levels in SST^+^ neurons was negated in these mice (**Fig.1B**, blue and red dots), suggesting that the elevated ER stress in SST^+^ neurons is mediated at least in part by PERK activity.

**Figure 1.**
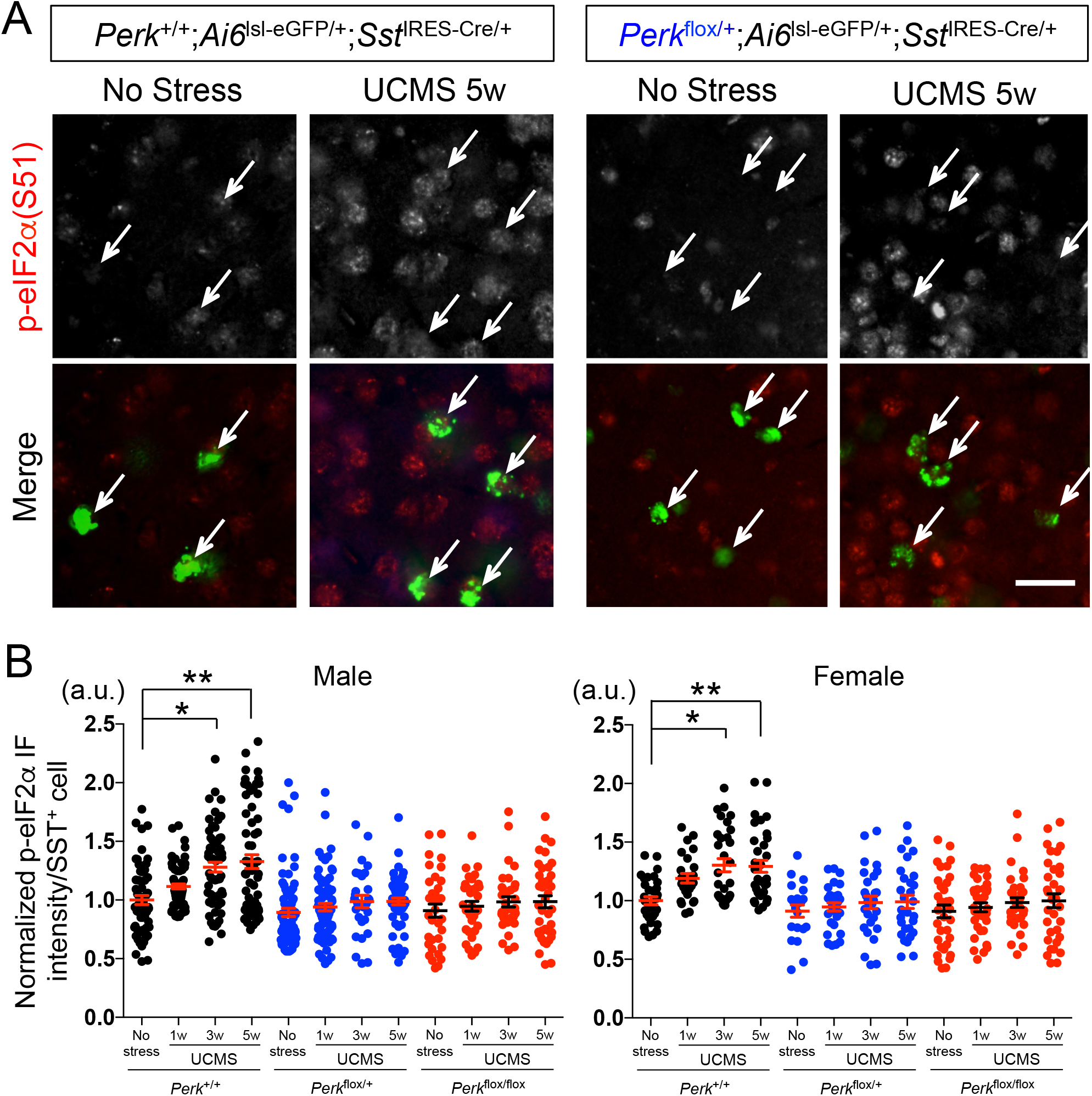
ER stress in SST neurons induced by UCMS. **A.** Phospho-eIF2α (Ser-51) levels (normalized by total eIF2α levels) in SST^+^ neurons in PFC of male control mice (*Perk*^+/+^;*Sst*^IRES-Cre^) as compared with mice with reduced *Perk* gene level (*Perk*^flox/+^;*Sst*^IRES-Cre^) before and after 5 weeks of UCMS. Arrow points to SST^+^ neurons labeled green (via *Ai6*^lsl-ZsGreen/+^;*Sst*^IRES-Cre/+^). Scale bar: 20 μm. **B.** Time-course of p-eIF2α levels in SST^+^ neurons during UCMS (1, 3 and 5 weeks). SST^+^ neurons in PFC of mice with reduced *Perk* levels show lower ER stress levels than those in control mice. (N=4/genotype/treatment/sex, 3-4 months old; Kruskal-Wallis test with Dunn’s multiple comparisons, *p<0.05, **p<0.01)

To next test the link between UCMS-induced ER stress and measures of behavioral emotionality, mice with varying *Perk* gene levels in SST^+^ neurons (i.e., *Perk*^+/+^; *Sst*^IRES-Cre/+^, *Perk*^flox/+^;*Sst*^IRES-Cre/+^ and *Perk*^flox/flox^;*Sst*^IRES-Cre/+^) were evaluated in a battery of assays for anxiety- and/or depressive-like behaviors before and after 5 weeks of UCMS (**Fig.2A**). Control mice with normal *Perk* gene levels exhibited UCMS-induced significant changes in a series of behaviors, such as decreased center time in open field test, decreased time in open arms of elevated plus maze, increased time in shelter zone in PhenoTyper, increased latency to feed in novelty-suppressed feeding test, decreased sucrose consumption, increased immobility time in forced swim test, deteriorated coat state (**Fig.S3**). When all 8 independent measures of emotionality behaviors are summarized by Z-scoring method (Guilloux et al., 2011), we confirmed significant effects of UCMS on elevated behavioral emotionality in all genotypes tested and in both sexes (**Fig.2B**). In contrast, mice with reduced *Perk* gene levels in SST^+^ neurons (hence suppressed ER stress levels, **Fig.1B**) showed significantly reduced behavioral emotionality scores (i.e. less anxiety-/depressive-like behaviors) in response to UCMS, compared to control mice exposed to UCMS (**Fig.2B** and **S3**). In male mice, *Perk* gene levels had no effect on behavioral emotionality at baseline (no stress conditions), whereas in female mice under no stress conditions, reducing *Perk* gene levels revealed decreased behavioral emotionality (**Fig.2B**), consistent with elevated baseline behavioral emotionality in females (Seney et al., 2018).

**Figure 2.**
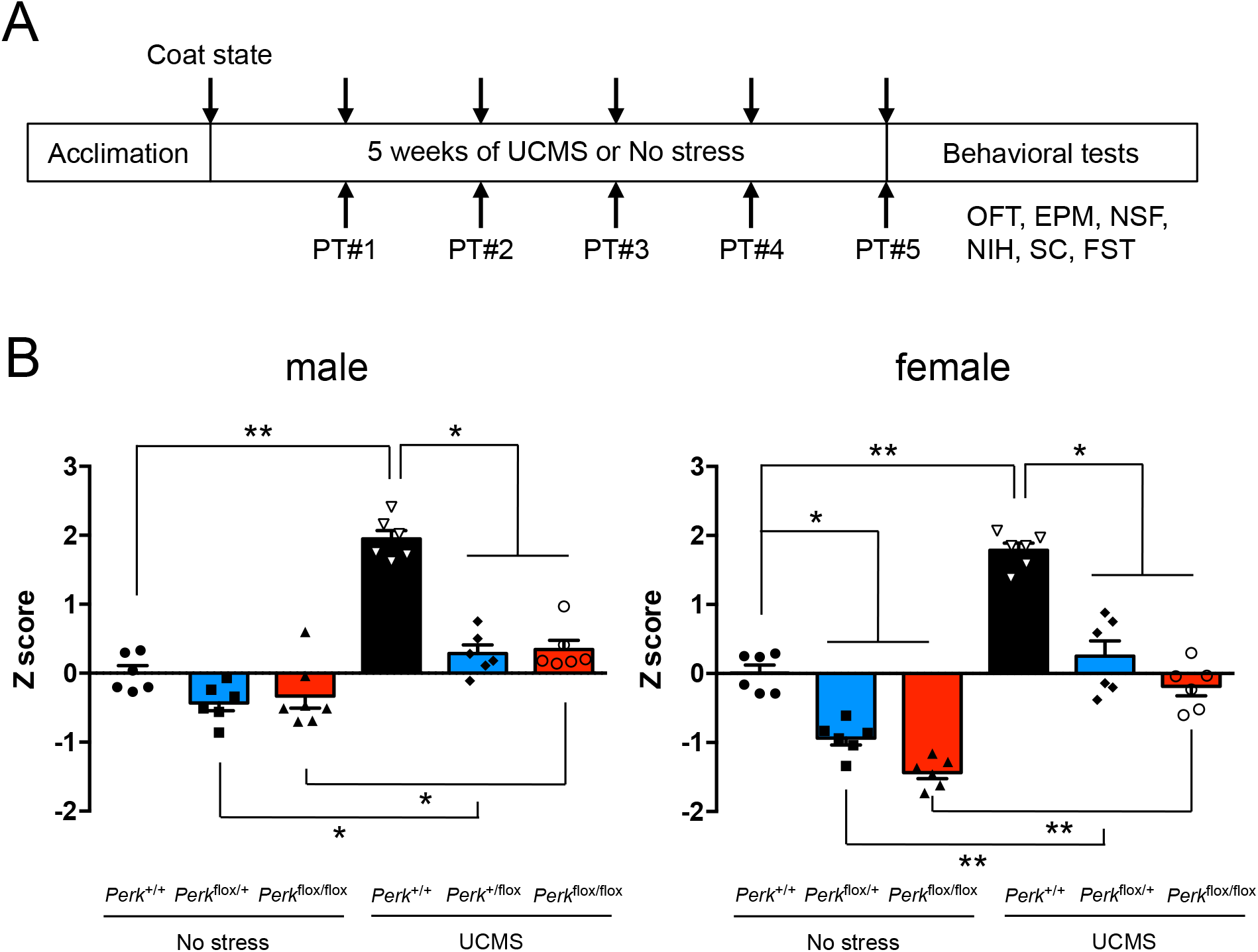
Genetic suppression of ER stress in SST^+^ neurons ameliorates UCMS-induced behavioral emotionality. **A.** Behavioral assay scheme: Mice (6♂+6♀/genotype, 3-4 months old) were subjected to 5 weeks of UCMS or kept under no stress, and tested for behavioral emotionality via 8 independent assays (weekly coat state evaluation and PhenoTyper (PT) followed by OFT, EPM, NSF, NIH, sucrose consumption (SC) and FST at the end of 5 weeks of UCMS). **B.** Z-scoring across all 8 assays shows consistent upregulated behavioral emotionality after UCMS in male and female mice. Blocking ER stress abolished UCMS-induced behaviors and lowered behavioral emotionality at baseline in female mice. Kruskal-Wallis test with Dunn’s multiple comparisons; *p<0.05, **p<0.01, ***p<0.001.

Although SST^+^ neurons show elevated ER stress upon UCMS, additional cells surrounding SST^+^ neurons also show increased phospho-eIF2α levels (**Fig.1A**). Given the ER stress-related transcriptomic changes found in pyramidal neurons (**Fig.S1A**), we tested a possible link between ER stress in pyramidal neurons and behavioral emotionality upon UCMS. Reducing *Perk* gene levels (hence suppression of ER stress) in CaMKII^+^ pyramidal neurons (i.e., *Perk*^flox/flox^;*Camk2*^Cre/+^, compared with *Perk^+/+^;Camk2^Cre/+^)* failed to rescue UCMS-induced behavioral emotionality (**Fig.S4**),

Together the data show that elevated ER stress in SST^+^ neurons, but not in CaMKII^+^ neurons, is sufficient to cause elevated behavioral emotionality.

### ER processing of preproSST induces elevated ER stress in SST^+^ neurons

To address the mechanistic basis for SST^+^ neuron-selective pathobiology, we focused on the SST^+^ neuron-intrinsic factor, SST precursor protein (preproSST), a polypeptide abundantly produced and processed through the ER to become a mature, bioactive SST peptide in SST^+^ neurons (Goodman et al., 1983). By analogy to pancreatic β cells, which show exacerbated ER stress through overproduction of insulin precursor protein (preproinsulin) under sustained hyperglycemic conditions (Arunagiri et al., 2018), we set out to test whether overexpression of preproSST causes elevated ER stress in SST^+^ neurons. Previous studies show that various acute stress modalities (e.g. cold shock, transient restraint, hypoxia) induce *Sst* mRNA expression in hypothalamus in rodents (Arancibia et al., 1984, 2000; Chen and Du, 2002; Priego et al., 2005; Polkowska and Wankowska, 2010). Similarly, we found that, during the initial phase of UCMS (3 days, 1 week), *Sst* mRNA levels are significantly elevated (~2-fold) in PFC compared to non-stressed control mice; however, the stress-induced increase in *Sst* mRNA expression became non-significant by 3 weeks of UCMS, and the net change in *Sst* levels became negative (~20% decrease) by 5 weeks of UCMS (**Fig.S5**). This suggests that the psychosocial stressors may initially increase the demand for SST production, but the capacity of SST^+^ neurons to produce SST deteriorates over time, leading to decreased SST expression as observed in brain disorders. A bi-phasic change in SST expression is also reported in hippocampus of epilepsy models; SST expression is acutely upregulated by seizures and reduced thereafter due to loss of SST^+^ neurons (Riekkinen and Pitkänen, 1990).

We next tested if forced expression of preproSST via stereotaxic adeno-associated viral (AAV) delivery, i.e., mimicking the increased psychosocial stress-induced SST production, was sufficient to induce elevated ER stress in PFC of mice. An AAV vector was designed to label cells green when infected in Cre-expressing cells (**Fig.3A**) and introduce preproSST into SST^+^ neurons (AAV-lsl-preproSST::2A::eGFP into *Sst*^IRES-Cre/+^ mice). This led to a marked increase in ER stress levels as compared to levels with GFP-only expression (**Fig.3B,C**). In contrast, expressing in SST^+^ neurons a cytosolic protein (e.g., PV, a marker of PV^+^ neurons) which does not need intracellular processing through ER or trans-Golgi network (AAV-lsl-PV::2A::eGFP into *Sst*^IRES-Cre/+^ mice) or introducing preproSST into a different cell type, CaMKII+ pyramidal neurons (AAV-lsl-preproSST::2A::eGFP into Tg:Camk2-Cre/+ mice), caused no apparent signs of elevated ER stress (**Fig.3C**). To directly test if an excessive need for intracellular processing of preproSST proteins may cause ER stress, a processing-incompetent preproSST mutant (lacking the signal peptide for ER insertion) or a mature form of SST (lacking proSST domain, thus no need of ER/Golgi processing and ready to be secreted) were expressed in SST^+^ neurons (**Fig.3A**). These mutants caused no apparent signs of elevated ER stress (**Fig.3C**).

**Figure 3.**
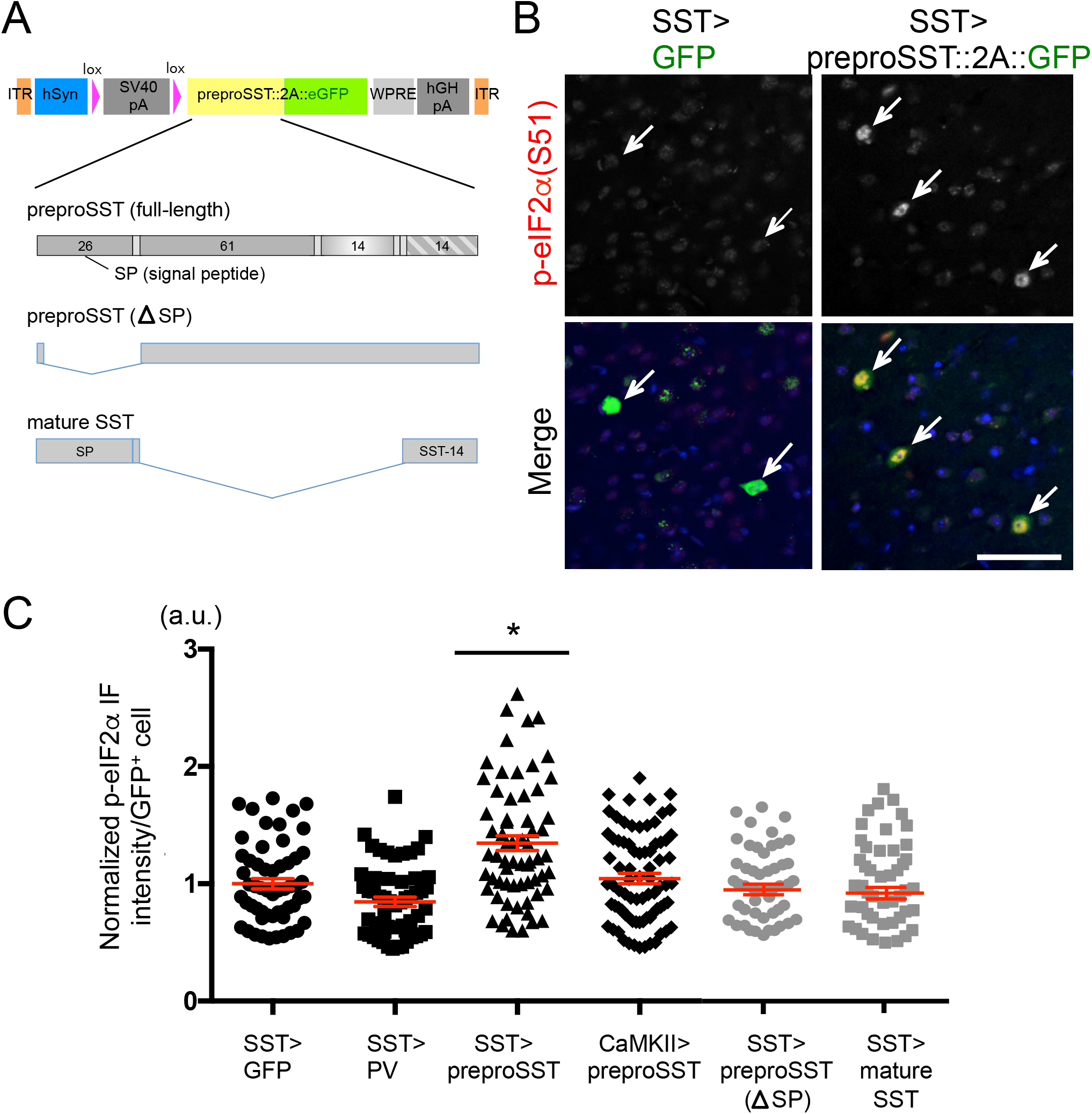
SST neuron-specific ER stress induced by preproSST. **A.** AAV-vector design: A human synapsin promoter (hSyn) drives the transgene expression when the host-derived Cre recombinase excises the stop cassette (lox-SV40 polyA-lox). Each transgene is fused in frame with eGFP via T2A self-cleavage signal. Transgene-expressing cells are labeled by eGFP, which is released from the expressed transgene product. Full-length preproSST and two truncation mutants (processing incompetent and mature SST) used are shown. **B.** GFP-tagged full-length preproSST transgene or GFP alone were introduced via AAV into SST^+^ neurons in PFC of male mice (N=4/group, 3-4 months old). The p-eIF2α (Ser-51) levels (normalized by total eIF2α levels) in GFP^+^ cells (pointed by arrow) were evaluated in IF assays. Scale bar: 50 μm. **C.** AAV encoding each transgene was delivered into a specific cell type in PFC (“cell type” > “transgene”) and the p-eIF2α levels in each infected cell were plotted (mean±SEM in red). Kruskal-Wallis test with Dunn’s multiple comparisons; *p<0.05.

We further tested the effect of preproSST-induced elevated ER stress on behaviors. Forced expression of preproSST in SST^+^ neurons caused elevated behavioural emotionality in mice (**Fig.S6**).

Together these results demonstrate that excessive preproSST processing is sufficient to induce ER stress and behavioral emotionality, suggesting it may be responsible for those phenotypes.

### UCMS-induced ER stress increases SST protein aggregation

SST has amyloid properties and co-aggregates with amyloid β in the brain of AD patients (Epelbaum et al., 2009; Solarski et al., 2018). Since elevated ER stress/UPR suggests the presence of misfolded or aggregated proteins in SST^+^ neurons, we tested whether UCMS could induce formation of aggregate-prone SST peptides in the PFC of mice. Using the filter-trap assay to capture insoluble protein species, we observed significantly increased SST aggregation in control mice (*Perk*^+/+^;*Sst*^IRES-Cre/+^) after 5 weeks of UCMS (**Fig.4A**). In contrast, levels of UCMS-induced SST aggregation were significantly suppressed in mice with reduced *Perk* levels in SST^+^ neurons (*Perk*^flox/flox^;*Sst*^IRES-Cre/+^) even after UCMS (**Fig.4A**), suggesting that SST aggregation occurs concomitantly with elevated ER stress in UCMS mouse brains. When the tissue extract was dissolved in RIPA buffer and analyzed by Western blot, detergent-soluble SST levels were either not significantly different across groups or showed a decreasing trend after UCMS (**Fig.4B**).

**Figure 4.**
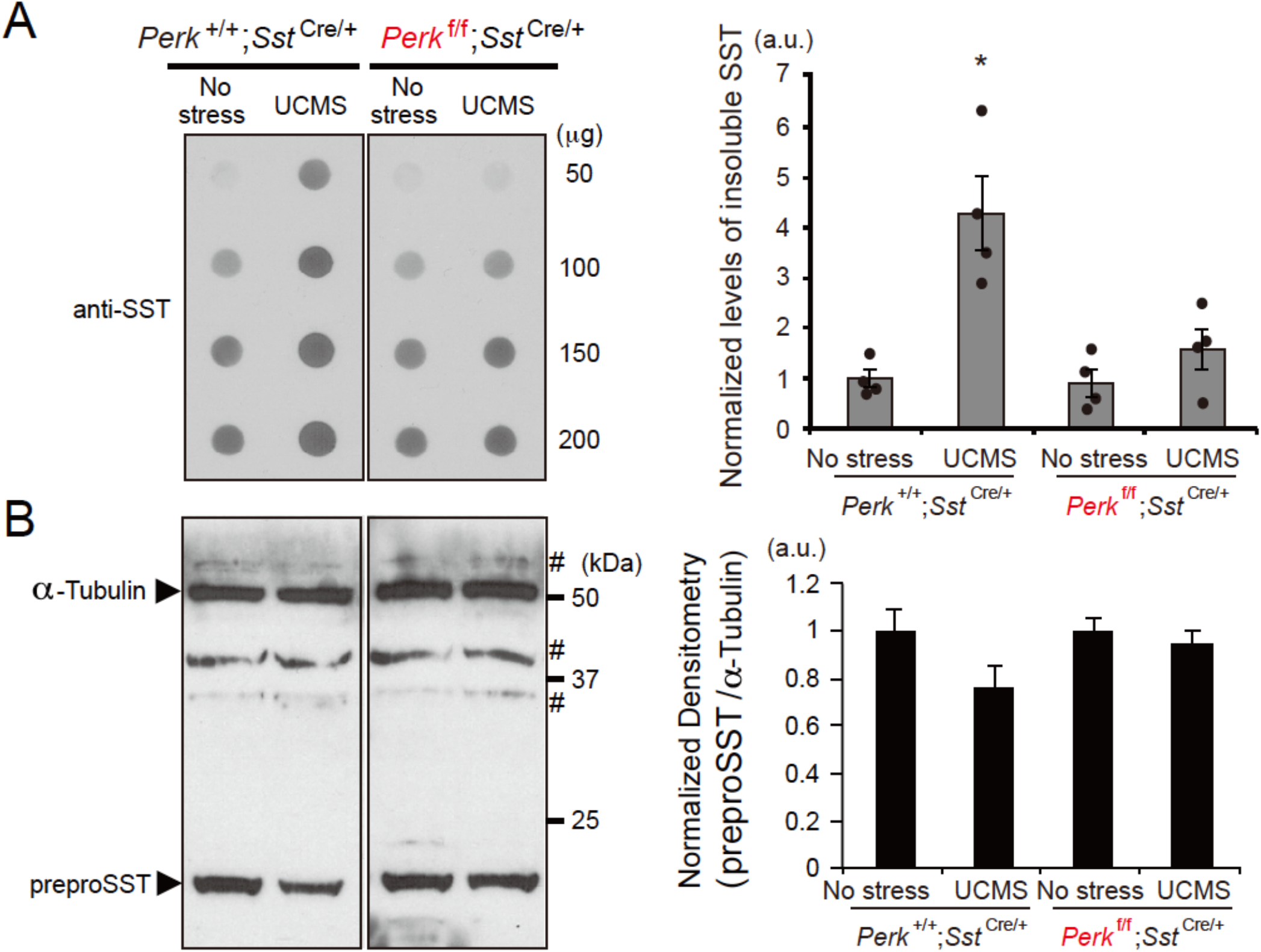
ER stress-dependent increase in insoluble preproSST peptides in PFC during UCMS. **A.** Proteins extracts (50, 100, 150 and 200 μg) prepared from PFC of mice (N=4/genotype/treatment, 3-4 months old, male) kept under no stress or 5 weeks of UCMS conditions were subjected to filter-trap assays to capture insoluble protein fractions and blotted with anti-SST antibody. The blot confirmed linearity of detection within the range of proteins used in this assay (50~200 μg). Graph shows the densitometry of insoluble SST proteins included in 100 μg of protein extracts from each sample. Kruskal-Wallis test with Dunn’s multiple comparisons; *p<0.05 (as compared with no stress *Perk^+I+^;Sst*^IRES-Cre^ control mice). **B.** Total SST peptides solubilized in 0.1% SDS were detected by Western blot. Each lane contains 25 μg protein extracts. α-Tubulin was used as loading control. # indicates non-specific immunoreactivity. Control mice (*Perk*^+/+^;*Sst*^IRES-Cre^) treated for 5 weeks of UCMS show a decreased trend of total soluble SST proteins (P=0.066, as compared with no stress *Perk^+I+^;Sst*^IRES-Cre^ control mice).

Together with findings from the previous section, these results suggest that sustained increased demand on preproSST processing leads to ER stress and aggregation of SST, representing a novel mechanism for the observed reduced SST in stress-related brain disorders.

## Discussion

The current study demonstrates that preproSST, the immature form of the SST neuropeptide, is an SST^+^ neuron intrinsic vulnerability factor, through the induction of intracellular ER stress, and that it could serve as a molecular origin for the selective vulnerability of SST^+^ neurons. When exposed to chronic environmental or psychosocial stressors, increased demand for SST production may exceed the physiological capacity of the ER/UPR system, leading to a vicious cycle of non-productive UPR overactivation and further disrupted proteostasis. This maladaptive response in the ER likely contributes to reduced SST availability and deregulated GABAergic neurotransmission in SST^+^ neurons as the underlying pathophysiological mechanism for stress-related brain disorders.

ER stress/UPR is primarily a cytoprotective mechanism responsible for surveillance and quality control of proteins. However, prolonged ER stress or aberrant UPR regulation leads to disrupted proteostasis through impaired processing in the ER and increased aggregate-prone protein species (Metcalf et al., 2020). Accumulation of aggregate-prone proteins is a hallmark of neurodegenerative disorders or aging (Hetz and Saxena, 2017), and is also beginning to be evident in neuropsychiatric conditions. The protein aggregates found in neurodegenerative disorders (e.g., Tau, amyloid-β, TDP-43, α-Synuclein and polyQ-containing Huntingtin) are generated either through aberrant posttranslational modifications (e.g., hyperphosphorylation, abnormal proteolytic cleavage) or missense and other genomic mutations, rendering the resulting proteins neurotoxic. By contrast, proteins that accumulate in the brains of psychiatric conditions may not be neurotoxic. Although accumulation or aggregations of several classes of proteins are reported in schizophrenia (e.g., p62 or ubiquitin^+^ protein species of unknown identity) (Sumitomo et al., 2018; Nucifora et al., 2019) or in autism (e.g., p62 and GABARAP) (Hui et al., 2019), these proteins have not been reported to be associated with neuronal cell death. In addition, a previous study reported increased levels of ER-resident chaperone proteins (i.e., GRP78/HSPA5/BiP, GRP94 and Calreticulin) in postmortem temporal cortex of MDD subjects who died by suicide compared with controls or MDD subjects who died by other reasons (Bown et al., 2000), yet the nature of aggregated proteins possibly accumulated in the brain of suicide/MDD subjects is unknown. A diverse array of functional proteins that are compromised through disrupted proteostasis may variably contribute to a constellation of symptoms in each disorder, depending on the nature and degree of aggregation as well as the cell type in which these aggregations occur.

In support of aggregate-prone nature of SST proteins, recent structural studies reported a striking feature of SST as a compactly folded protein with an amyloid-like structural property when it is packaged into secretory vesicles (Solarski et al., 2018). SST is also reported to physically interact with amyloid-β and co-sediments with amyloid plaques in postmortem AD brains (Wang et al., 2017). Currently, while we showed increased aggregation of SST proteins in UCMS brains, we do not know the nature of aggregate-prone protein species in UCMS mice, in particular, whether additional proteins may co-aggregate with SST proteins or whether aggregated SST proteins serve as seed for additional proteins to aggregate. Nonetheless, reduced levels of functionally available SST proteins through disrupted proteostasis in UCMS mouse model provide a testable hypothesis that SST proteins may be compromised through a similar mechanism in MDD brains. Notably, overexpressed preproSST caused elevated ER stress in SST^+^ neurons, but not in pyramidal neurons. This suggests that pyramidal neurons may have a greater UPR capacity than SST^+^ neurons, or that SST^+^ neuron-intrinsic factors may render this cell type vulnerable to disrupted ER homeostasis.

A recent study demonstrated the role of both pyramidal and SST^+^ neurons in long-term memory consolidation via eIF2α-dependent protein translation in hippocampus (Sharma et al., 2020). In contrast, the current study demonstrates the predominant role of SST^+^ neurons in emotionality control via PERK/eIF2α signalling under prolonged stress conditions. Given the unique role of SST^+^ neurons in gating information processing toward pyramidal neuron dendrites within the corticolimbic microcircuitry (Prévot and Sibille, 2021), it is possible that the cognitive processing is mediated by the coordinated activity of both neuron types. In this regard, it is tempting to speculate that chronic stress-induced cognitive deficits may be rescued by enhancing eIF2α signalling in pyramidal neurons or in both pyramidal and SST^+^ neurons. Additional experiments are necessary to further dissect the circuit mechanisms for emotion versus cognition and to determine the extent of convergence or independence of multiple cell types regulating these two behavioral dimensions.

While our study demonstrates the mechanism for elevated ER stress in emotionality disturbances, analogous mechanisms for elevated ER stress may be shared by other, distinct pathophysiological contexts. In diabetes, ER stress is exacerbated specifically in pancreatic β cells by increased demand for insulin production and secretion under the hyperglycemic conditions (Arunagiri et al., 2018). Notably, a series of point mutations in the *preproinsulin* gene compromise intracellular processing through the ER and trans-Golgi network, leading to accelerated onset of diabetes (Liu et al., 2015). Elevated ER stress is also implicated in the pathophysiology of neurodegenerative disorders, including AD and prion disease (Hetz and Saxena, 2017). Accumulation of intracellular aggregates (i.e., neurofibrillary tangles, Tau) in affected neurons activates the UPR pathway in the ER lumen via activation of PERK, resulting in compromised neuronal function and cell death. Suppressing ER stress by administration of a PERK inhibitor could ameliorate neuronal apoptotic events in these AD models (Rozpedek et al., 2015; Gerakis and Hetz, 2018). These results suggest that cell type-specific vulnerability to ER stress may represent a yet-unrecognized family of diseases with a common pathophysiological cellular mechanism, although the organs and cell types affected may differ across diseases.

## ACKNOWLEDGMENTS

The authors thank Mohan Pabba, Rammohan Shukla, Mounira Banasr, Thomas Prevot, and Hyunjung Oh for comments or discussion. This work was supported by grants from the Canadian Institute of Health Research (CIHR #153175 to ES), National Alliance for Research on Schizophrenia and Depression (NARSAD award #25637 to ES), the National Institutes of Health (MH-093723 to ES), Campbell Family Mental Health Research Institute (to ES). The authors declare no competing interests.

## AUTHOR CONTRIBUTIONS

TT and ES conceived the study and designed the experiments; TT and AS acquired and analyzed data; DN analyzed gene expression profiles; TT and ES wrote and edited the manuscript.

## Experimental procedures

### Transcriptomic analysis

Twenty male C57Bl/6J mice (JAX stock #000664) were exposed to either control or UCMS conditions (n=10/group) for 5 weeks and underwent a battery of anxiety- and anhedonia-like behavioral tests before sacrifice, as described (Nikolova et al., 2018). Pyramidal, SST^+^, PV^+^ and VIP^+^ neurons (130 somas/cell-type/mouse) were collected from the mPFC using fluorescent in situ hybridization and laser capture microdissection as described (Rocco et al., 2017). RNAseq was performed as described (Shukla et al., 2019), using the HiSeq 2500 platform (Illumina, San Diego, CA). Gene-set enrichment analysis (GSEA) was used to assess UCMS-induced transcriptomic changes in each cell type (Subramanian et al., 2005).

### Animals

Conditional *Perk* knockout mice (JAX stock #023066) and transgenic mice that express Cre in *Sst* locus (JAX stock #013044), Camk2-Cre (JAX stock #005359) or ZsGreen (JAX stock #007906) were obtained from The Jackson Laboratory (Bar Harbor, ME, USA). Mice were maintained on the C57BL/6J genetic background for at least 10 generations. Two to 4 months old male mice were used for experiments. All animal assays and data collection were performed by experimenters who were blind to animal genotypes. Maintenance and use of animals were in accordance with the NIH Guide for the Care and Use of Laboratory Animals, and approved by the Institutional Animal Care and Use Committees at CAMH.

### Stereotaxic injection of adeno-associated virus (AAV)

Adeno-associated virus (AAV) (~5×10^12^ gc/ml) encoding murine preproSST transgene fused with GFP via T2A self-cleavage spacer sequence was prepared by AAVpro purification kit (TaKaRa Bio, Japan) and bilaterally injected into PFC (AP +1.9 mm; ML ±0.15 mm; DV −0.8 mm), using a stereotaxic apparatus (RWD Life Science, USA). A total of 0.5 μl of purified virus was delivered on each hemisphere over a 5-min period using glass microcapillary pipettes (1.14 mm O.D. x 3.5” length x 0.53mm I.D.) and the Nanoject II injector (Drummond, PA, USA). Mice (8 to 10 weeks of age) were anesthetized by isoflurane inhalation during the stereotaxic injection, and used for downstream assays 3 weeks post-injection.

### Behavioral assays

After one-week acclimation to the test environment, mice were subjected to either non-stress or UCMS conditions for 5 weeks, and the coat state and the shelter zone duration in PhenoTyper (Noldus) (post-light duration in the shelter zone) were evaluated weekly. Mice were then tested in a battery of anxiety- and anhedonia-like behavioral assays in the following order: open field test (OPT, center time during the first 5 min of tests), elevated plus maze (EPM, %Time in open-arm, frequency of entry to open arm), novelty-induced hypophagia (NIH, latency to feed), rate of sucrose consumption (24h) (normalized by water consumption during 24h), novelty-suppressed feeding (NSF, latency to feed) and forced swim test (FST, %Immobility time), in order to minimize potential influences of the preceding tests on subsequent assays.

### Quantitative reverse transcription-polymerase chain reaction (qRT-PCR)

Total RNA was extracted from PFC using RNeasy Mini kit (Qiagen, USA), and reverse-transcribed with a ReverTra Ace cDNA synthesis kit (Toyobo, Japan). TaqMan probes were purchased from Applied Biosystems, Inc. (ABI, USA). All data were normalized with *Gapdh* as reference.

### Filter-trap assay and Western blot

Filter trap assays were performed as described (van Waarde-Verhagen and Kampinga, 2018). In brief, brain samples were prepared in PBS containing 1% sodium dodecyl sulfate, and blotted on the cellulose acetate membrane using a Bio-Dot apparatus (BioRad, USA). The membrane was then probed with anti-SST (1:200, rabbit polyclonal, H-106, Santa Cruz Biotechnology, USA) and HRP-conjugated secondary antibody, and the trapped proteins were detected using the ECL kit and ChemiDoc imaging system (BioRad). For detection of total proteins in brain extracts, standard Western blot was performed using anti-SST and anti-α-Tubulin (1:8,000, B-5-1-2, mouse monoclonal, Sigma).

### Immunofluorescence analysis

For immunofluorescence analysis of tissue sections, brains of perfused mice (*n=4* per group) were serially cut into 35 μm-thick coronal sections using cryostat (Leica, Germany). Sections were permeabilized in PBS containing 0.3% Triton X-100 for 30 min, incubated for 30 min at room temperature in 10% goat serum (Chemicon) in PBS, and then immunostained for 16 h at 4°C with primary antibodies followed by Alexa Fluor^®^ 568- or 633-conjugated secondary antibodies (Molecular Probes) for 1 h at room temperature. Antobodies used: anti-eIF2α (1:200, mouse monoclonal, Abcam, ab5369), anti-phospho-eIF2α (Ser-51) (1:200, rabbit monoclonal, Cell Signaling Technology #3398). The stained samples were observed using IX81 microscope equipped with the Disc-Spinning Unit (Olympus, 40x objective lens, NA=1.3); images of one optical section (1 μm thick) were acquired from 3 to 6 non-overlapping areas per section, randomly chosen in PFC region, and 3 to 4 serial sections were analyzed. The fluorescence intensities of immunostaining were measured from each neuronal soma (30~50 somas per section) using ImageJ (NIH), with the background levels of staining in adjacent regions being subtracted, and the normalized average immunofluorescence intensity (phospho-eIF2α/eIF2α) was calculated across all serial sections from every mouse used.

### Statistical Analysis

Sample size for each animal experiment was predetermined to ensure adequate statistical power for drawing conclusion. Statistical analysis was performed using Prism software (GraphPad). Behavioral and molecular experiments were statistically analyzed either by repeated measures two-way analysis of variance (ANOVA) followed by Bonferroni post-hoc tests or Kruskal-Wallis test followed by Dunn’s multiple comparisons, unless otherwise stated in the figure legend. P < 0.05 was considered statistically significant.

