## Supplementary Figure Legends for "Molecular origin of somatostatin-positive neuron vulnerability"

By

Toshifumi Tomoda^1*^, Akiko Sumitomo^1^, Dwight Newton^1^, and Etienne Sibille^1,2,3,*^

^1^Campbell Family Mental Health Research Institute, Centre for Addiction and Mental Health (CAMH), Toronto, Ontario M5T 1R8, Canada

^2^Department of Psychiatry, University of Toronto, Toronto, Ontario M5T 1R8, Canada

^3^Department of Pharmacology and Toxicology, University of Toronto, Toronto, Ontario M5T 1R8, Canada

**Supplementary Figures 1-6 Legends**

**Supplementary Fig. 1. Gene set enrichment analysis of ER stress/UPR-related pathways expressed in neurons of cortical circuitry in UCMS mice.**

**A.** Gene set enrichment analysis (GSEA) summary of ER stress/UPR-related pathways significantly enriched in neurons of cortical circuitry in UCMS mice. PYR: pyramidal neurons; SST: SST^+^ neurons; NES: normalized enrichment score.

**B.** Enrichment plot for each pathway shown in **A**.

**Supplementary Fig. 2. Gene set enrichment analysis of three regulatory pathways for ER stress/UPR expressed in cortical SST^+^ neurons of UCMS mice.**

**A.** GSEA summary of PERK, IRE1α and ATF6 pathways expressed in cortical SST^+^ neurons of UCMS mice.

**B.** Enrichment plot for each pathway shown in **A**.

**C.** Heatmap analysis of selected representative genes in each pathway. Gene name and normalized enrichment score are shown in each box.

**Supplementary Fig. 3. Genetic suppression of ER stress in SST^+^ neurons ameliorates UCMS-induced behavioral emotionality in mice.**

Mice (6♂+6♀/genotype, 3–4 months old) were subjected to 5 weeks of UCMS or kept under no stress (NS) conditions, and assayed for open field test (OPT), elevated plus maze (EPM), PhenoTyper (PT), novelty-suppressed feeding (NSF), novelty-induced hypophagia (NIH), sucrose consumption (SC), forced-swim test (FST) and coat state. Statistical significance was evaluated by two-way ANOVA with repeated measures, followed by Bonferroni post-hoc tests; *P<0.05, **P<0.01 (as compared to the control genotype (*Perk*^+/+^;*Sst*^IRES-Cre/+^) of the same treatment group (i.e., either NS or 5 weeks of UCMS)); ^#^P<0.05, ^##^P<0.01, ^###^P<0.001 (as compared to the same genotype of the NS group).

**Supplementary Fig. 4. No rescue of UCMS-induced behavioral emotionality in mice with genetic suppression of ER stress in CaMKII^+^ pyramidal neurons.**

Mice (6♂+6♀/genotype, 3–4 months old) were subjected to 5 weeks of UCMS or kept under no stress, and tested for behavioral emotionality. Z-scoring of behavioral emotionality as integral summary of all 8 assays are shown. Kruskal-Wallis test with Dunn’s multiple comparisons; **P<0.01.

**Supplementary Fig. 5. *Sst* mRNA levels in wild-type mice during 5 weeks of chronic stress.**

Mice (N=6/group, 3-4 months old male) were subjected to chronic stress (physical restraint in a Falcon tube for 1 h, twice per day) for indicated periods and the PFC was sampled for quantitative PCR analysis immediately (15 min) after the last stress procedure. Data are plotted as mean±SEM. Kruskal-Wallis test with Dunn’s multiple comparisons; *P<0.05 (as compared with the no stress condition).

**Supplementary Fig. 6. Forced expression of preproSST in SST^+^ neurons induces behavioral emotionality.**

Mice (*Sst*^IRES-Cre/+^, 6♂+6♀/group, 3 months old) were stereotaxically injected into PFC with either AAV-lsl-GFP control or AAV-lsl-preproSST::T2A::GFP virus, and tested for behavioral emotionality in OFT and EPM assays 3 weeks post-surgery. The effects of preproSST transgene expression on both OFT and EPM were significant when all cohorts (males and females combined) were evaluated (P=0.046 for OFT, P=0.048 or EPM). When the effects were evaluated for each sex, OFT in males (P=0.040) and EPM in females (P=0.042) showed significant effects. Statistical significance was evaluated by Mann-Whitney U test (*P<0.05, as compared with control AAV of the same sex).
