## Supplementary Figures for "Molecular origin of somatostatin-positive neuron vulnerability"

### Supplementary Fig. 1

A

ER stress/UPR-related pathways enriched in neurons of cortical circuitry in UCMS mice

| Cell Type | Pathway | ONTOLOGY ID | NES | NOM p-val |
|---|---|---|---|---|
| PYR | CELLULAR RESPONSE TO HEAT | GO:0034605 | 1.6070465 | 0.011447419 |
| PYR | RESPONSE TO HEAT | GO:0009408 | 1.5792518 | 0.006862371 |
| SST | UBIQUITIN-DEPENDENT ERAD PATHWAY | GO:0030433 | 1.4322895 | 0.03341784 |
| SST | ERAD PATHWAY | GO:0036503 | 1.3694564 | 0.042072736 |
| SST | CHAPERONE-MEDIATED PROTEIN FOLDING | GO:0061077 | 1.7146109 | 0.002917415 |
| SST | CHAPERONE COFACTOR-DEPENDENT PROTEIN REFOLDING | GO:0051085 | 1.670395 | 0.013686534 |
| SST | 'DE NOVO' PROTEIN FOLDING | GO:0006458 | 1.5231644 | 0.027027028 |
| SST | 'DE NOVO' POSTTRANSLATIONAL PROTEIN FOLDING | GO:0051084 | 1.5152692 | 0.035882484 |
| SST | RESPONSE TO HEAT | GO:0009408 | 1.4188943 | 0.034334764 |
| SST | HEAT SHOCK PROTEIN BINDING | GO:0031072 | 1.414919 | 0.020537898 |

B

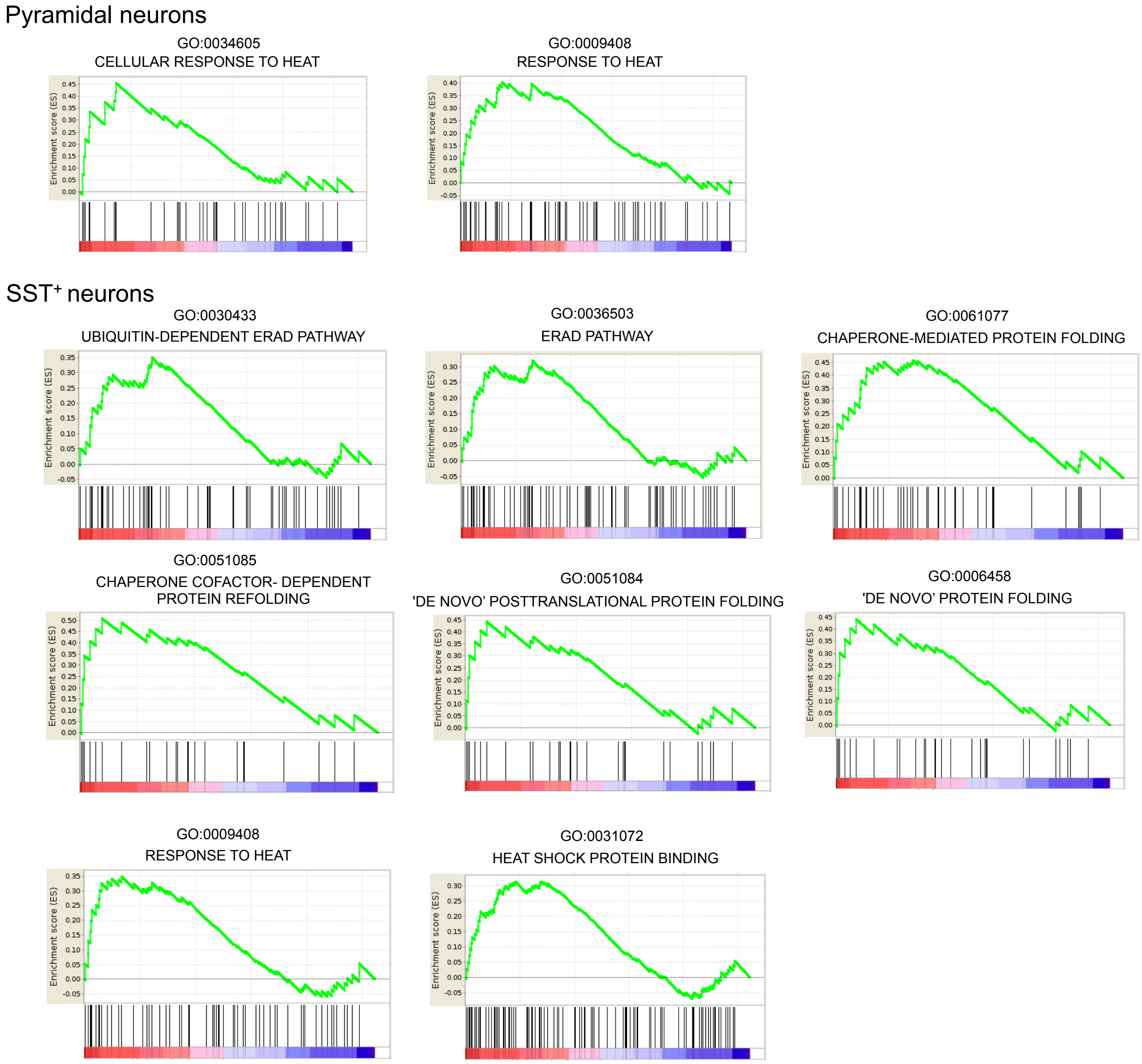

Supplementary Fig. 2

A

| Pathway | ONTOLOGY ID | NES | NOM p-val |
|---|---|---|---|
| PERK pathway genes | REACTOME_381042 | 1.404323 | 0.079627 |
| IRE1α pathway genes | GO:0036498 | -1.23048 | 0.145998 |
| ATF6 pathway genes | GO:0036500 | 0.704001 | 0.836375 |

B

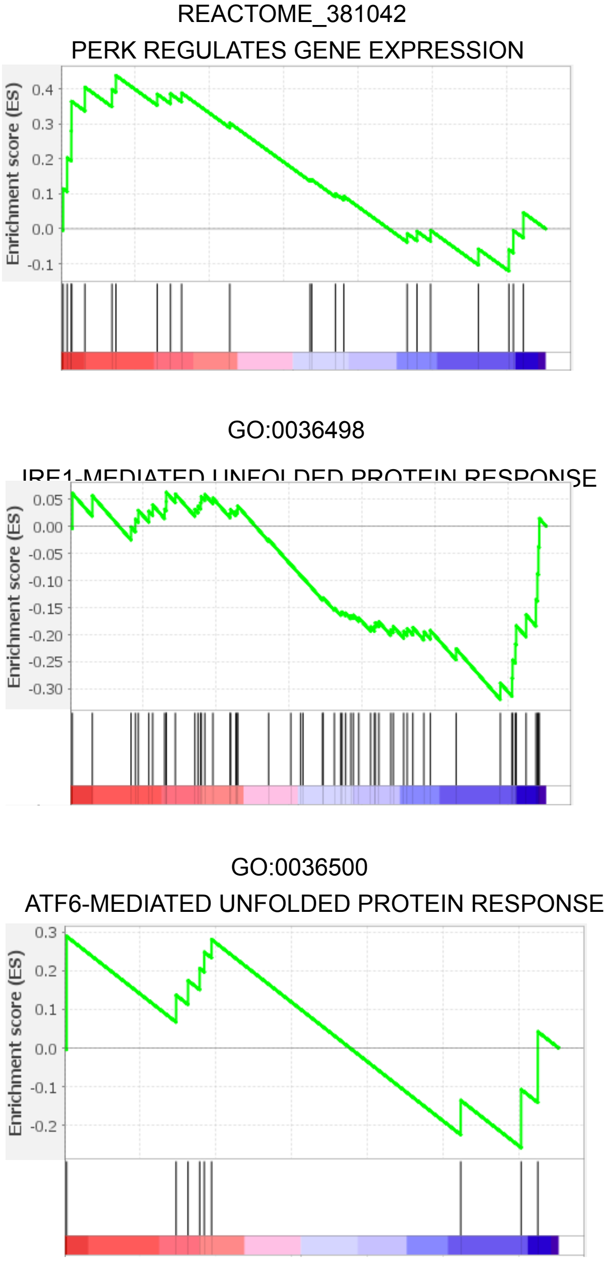


C

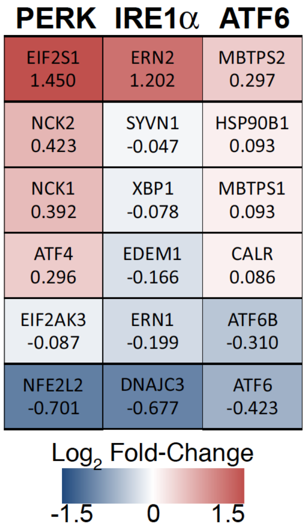

### Supplementary Fig. 3

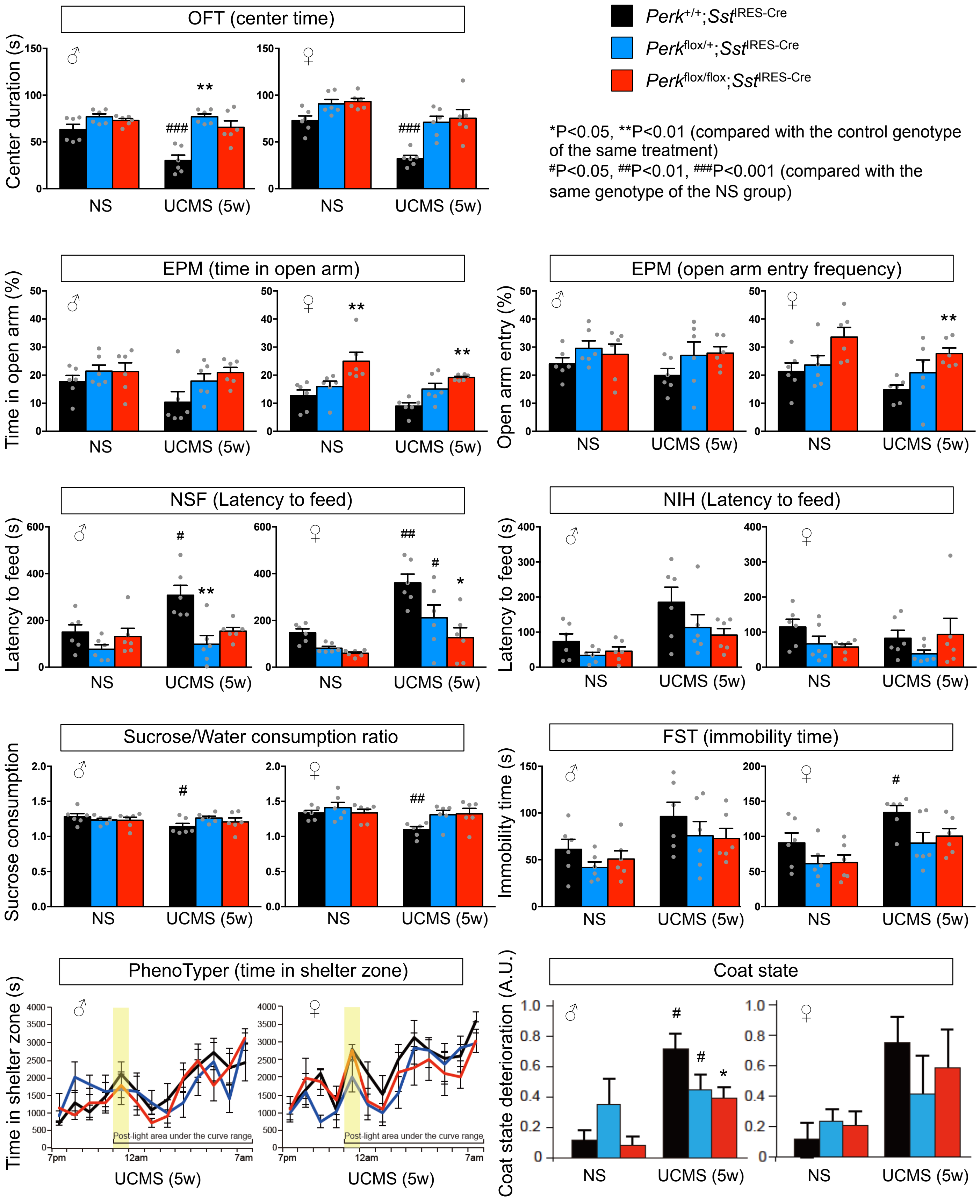

### Supplementary Fig. 4

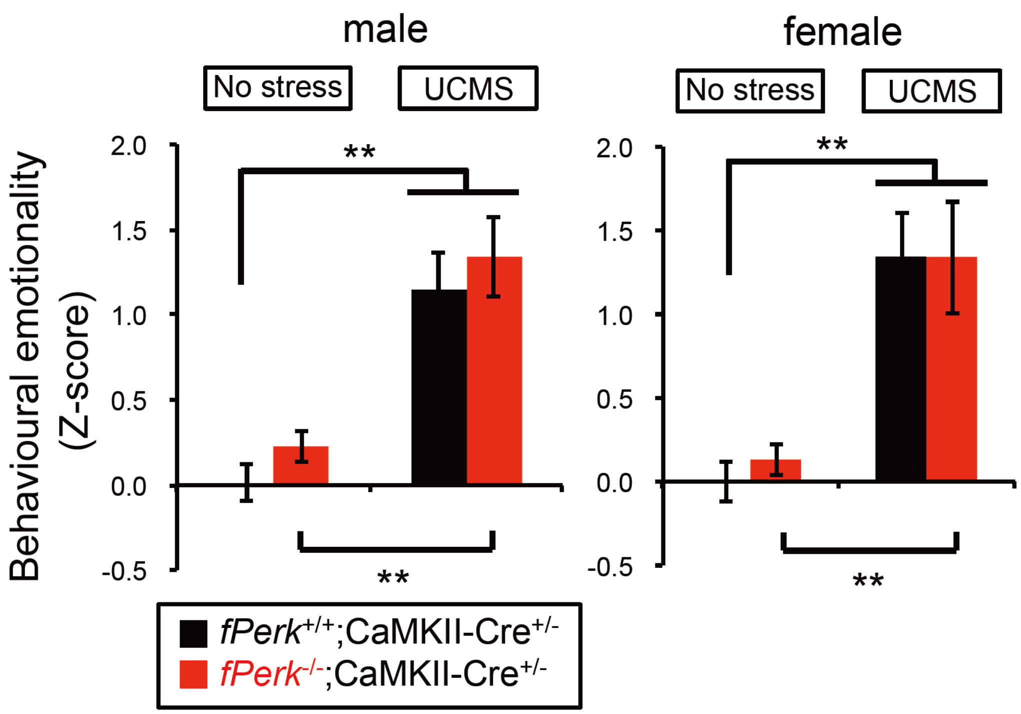

Supplementary Fig. 5

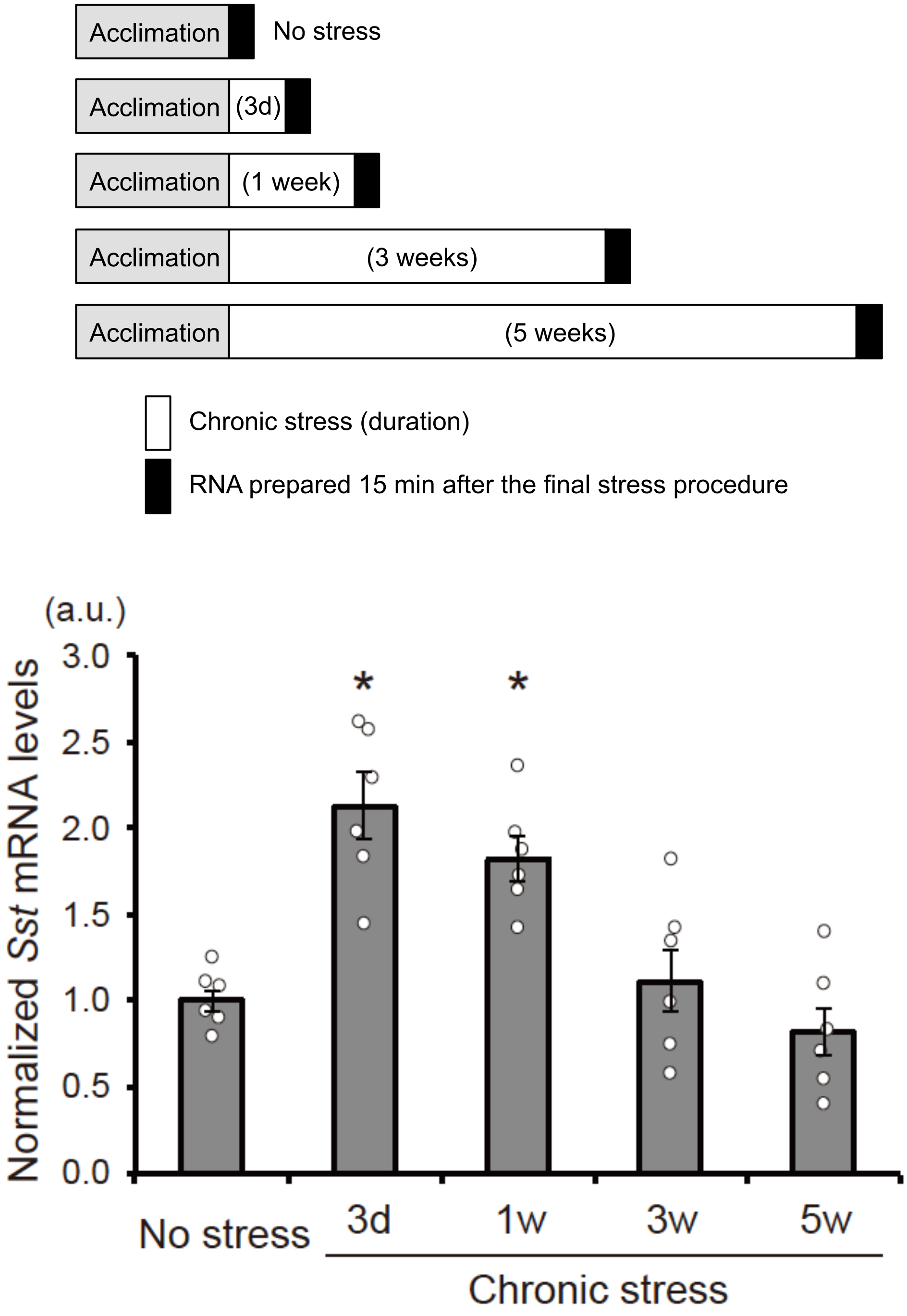

### Supplementary Fig. 6

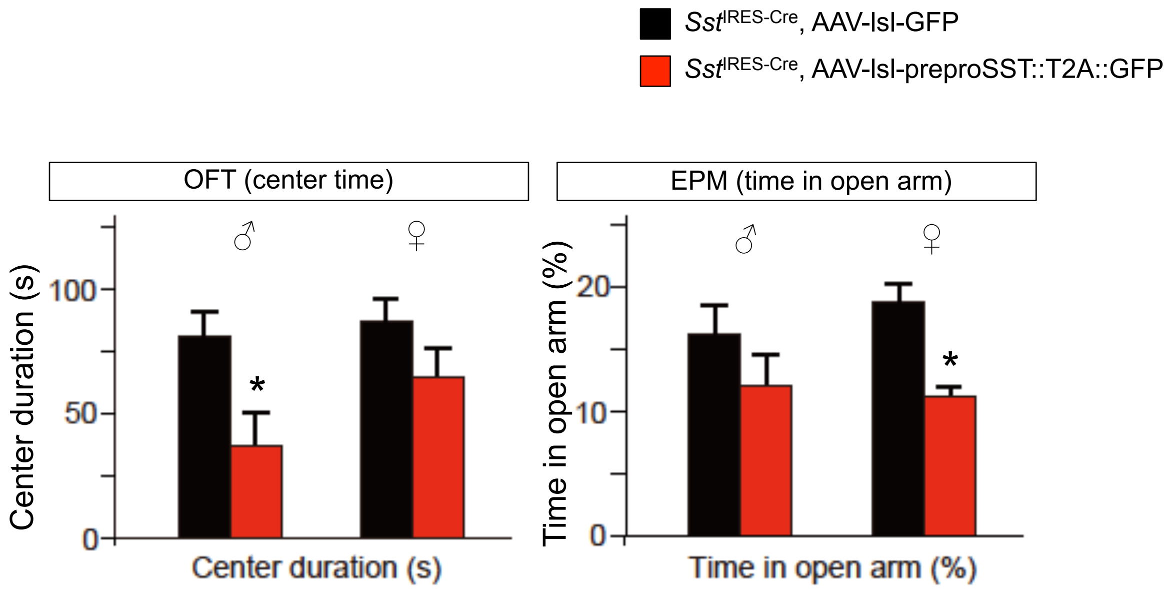
